# Genetic regulation of microRNAs in the older adult brain and their contribution to neuropsychiatric conditions

**DOI:** 10.1101/2024.09.10.610174

**Authors:** Selina M. Vattathil, Ekaterina S. Gerasimov, Se Min Canon, Adriana Lori, Sarah Sze Min Tan, Paul J. Kim, Yue Liu, Eric C. Lai, David A. Bennett, Thomas S. Wingo, Aliza P. Wingo

## Abstract

MicroRNAs are essential post-transcriptional regulators of gene expression and involved in many biological processes; however, our understanding of their genetic regulation and role in brain illnesses is limited. Here, we mapped brain microRNA expression quantitative trait loci (miR-QTLs) using genome-wide small RNA sequencing profiles from dorsolateral prefrontal cortex (dlPFC) samples of 604 older adult donors of European ancestry. miR-QTLs were identified for 224 miRNAs (48% of 470 tested miRNAs) at false discovery rate < 1%. We found that miR-QTLs were enriched in brain promoters and enhancers, and that intragenic miRNAs often did not share QTLs with their host gene. Additionally, we integrated the brain miR-QTLs with results from 16 GWAS of psychiatric and neurodegenerative diseases using multiple independent integration approaches and identified four miRNAs that contribute to the pathogenesis of bipolar disorder, major depression, post-traumatic stress disorder, schizophrenia, and Parkinson’s disease. This study provides novel insights into the contribution of miRNAs to the complex biological networks that link genetic variation to disease.

## 1 Introduction

MicroRNAs (miRNAs) are small non-coding RNAs that suppress gene expression by binding complementary mRNA sequences and causing either transcript degradation or translation repression. miRNAs are typically produced through multi-step post-transcriptional processing that begins with cleavage of the primary miRNA transcript (hundreds or thousands of nucleotides long) into a hairpin-shaped precursor miRNA (pre-miRNA), which is transported into the cytoplasm and then cleaved into ∼22 nucleotide mature miRNAs^1^. Each pre-miRNA can produce up to two mature miRNAs, one from each arm of the hairpin. Each miRNA can influence the expression of potentially hundreds of genes, and thereby exert a broad influence on gene expression^2^, which make them relevant for understanding the complex networks of interacting genes involved in most human disease.

While miRNAs have a well-described role in gene regulation, less is known about how they themselves are genetically regulated. By contrast, comparatively more is known about how genetic variants are associated with variation in transcript and protein expression – so called quantitative trait loci (QTL) mapping. This has led to important insight into human conditions, especially by integrating QTLs with genome-wide association study (GWAS) results^3–6^. Thus, we hypothesize that miRNA QTLs (miR-QTLs) ought to provide similar insights.

Characterizing miR-QTLs has several additional benefits. First, since miRNAs regulate genes that may be distally located in the genome, miR-QTLs could be used to reduce the multiple testing burden for finding trans-pQTLs. Second, miR-QTLs could be used to test whether miRNAs mediate protein expression when both the miRNA and protein are under genetic control, which would complement existing miRNA target prediction approaches. Together, these results would provide a more complete picture of gene expression regulation.

Given their small size, miRNAs identified from next-generation sequencing data are prone to false annotation if appropriate criteria are not applied^7,8^. Coupled with the fact that miRNAs may be cell type-specific^9^, the precise complement of human miRNAs remains a subject of debate^10,11^.

Here, we first cataloged miRNA QTLs (miR-QTLs) using small RNA sequencing from adult human dorsolateral prefrontal cortex (dlPFC) (Figure 1). Given the current uncertainty about the true miRNA complement, we conducted parallel miR-QTL mapping using miRNAs from two miRNA databases that have different inclusion criteria. Next, we leveraged the high confidence miR-QTLs to address questions about gene regulation, such as whether miRNAs are regulated independently of host genes; whether miR-QTLs are enriched in promoters or enhancers; whether miR-QTLs can be used to detect trans-pQTLs and support predicted miRNA targets; and whether miR-QTLs are tissue-specific. Finally, we applied multiple complementary methods for integrating the miR-QTLs with GWAS signals from 16 neurodegenerative and psychiatric traits to identify miRNAs that mediate the effect of genetic variation on the brain-related traits (Figure 1). Altogether, we uncovered miRNAs involved in brain illnesses, characterized features of miRNA regulation, and provide a human brain miR-QTL resource for future study.

**Figure 1.**
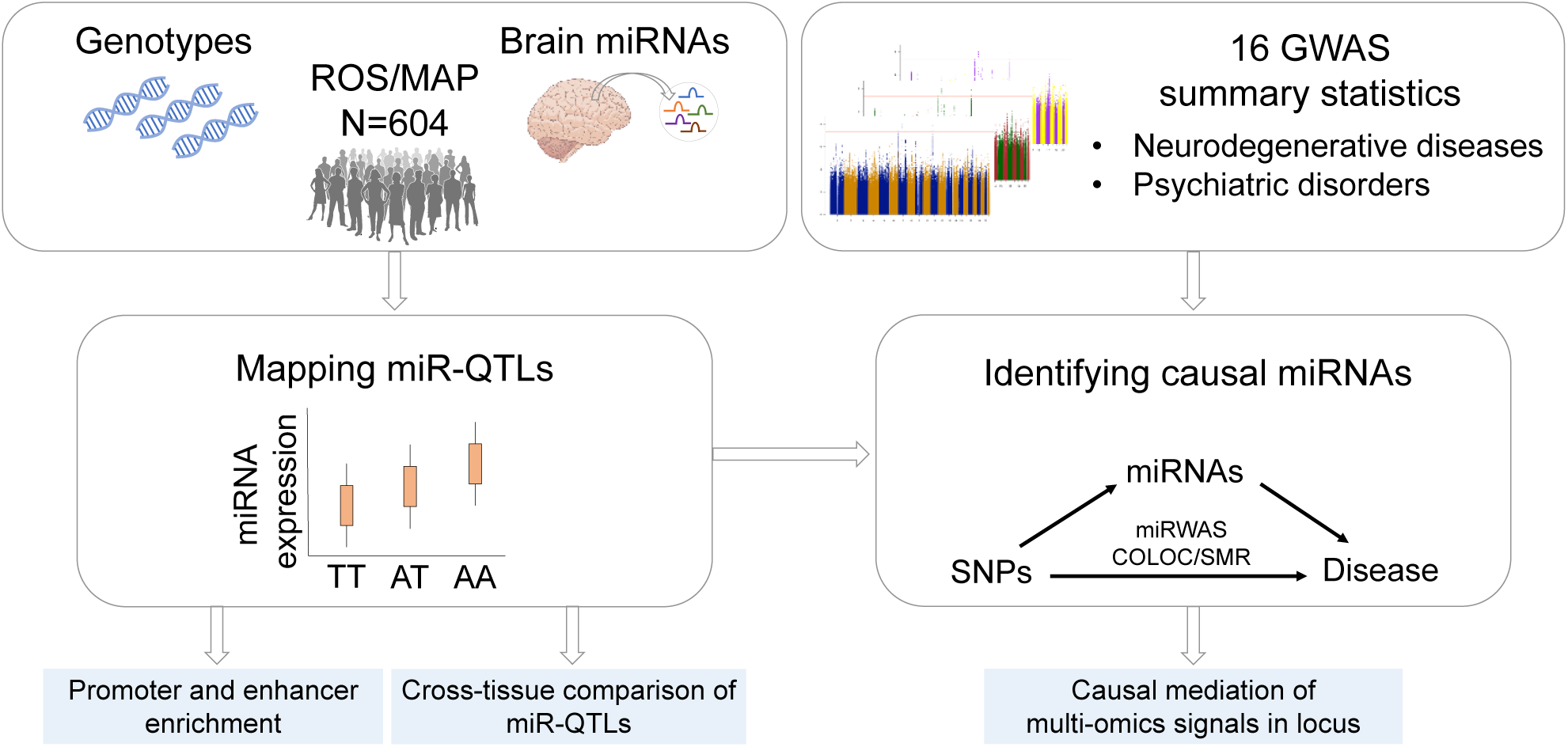
Main analyses.

## 2 Results

### 2.1 miR-QTL mapping

miRNA and genotype data from 604 individuals were available for analysis after quality control filtering. Sample and data characteristics are summarized in Supplementary Table 1. All participants were over 70 years old at the time of death, and the median age of death was 90 years. The median miRNA sequencing depth per sample was 27.3 million reads (range 13.2 million to 37.4 million). We chose to independently map miRNA sequencing reads to two reference miRNA databases – miRBase^12^ and MirGeneDB^13,14^. The rationale for this was that miRBase is a widely used source that is likely valuable to the research community. Yet, miRBase has been noted for its substantial number of falsely annotated miRNAs^8,15–17^. On the other hand, MirGeneDB is a manually curated miRNA database with reported low rate of false annotations and high completeness for human miRNAs^13,14^.

Using miRBase to map miRNA reads resulted in a median of 18.8 million mapped reads per sample (range 7.7 million to 27.4 million) (Supplementary Figure 1). After filtering based on our abundance criteria (≥ 1 RPM for ≥ 50% of samples), there were 624 autosomal miRNAs available for miR-QTL analysis (Supplementary Tables 2-3). Considering SNPs within 500kb of the precursor sequences, we tested 1,315,257 SNP-miRNA pairs and observed miR-QTLs for 326 miRNAs at false discovery rate (FDR) < 1% (unadjusted P-value < 2.98 × 10^-4^). Hereafter, miRNAs with at least one miR-QTL are referred to as eMiRs. Since we analyzed a dense set of genotypes, many SNPs are in high linkage disequilibrium (LD). To identify a reduced set of SNPs, we applied LD-based clumping to the miR-QTL results to identify ‘index’ miR-QTLs for each eMiR, followed by conditional analysis to identify conditionally independent miR-QTLs for each eMiR. Of the 326 eMiRs, 71% had one conditionally independent miR-QTL, 22% had 2, and 7% had three, four, or five.

Using MirGeneDB to map miRNA reads resulted in a median of 18.7 million mapped reads per sample (range 7.7 million to 27.3 million). The list of tested MirGeneDB miRNAs and RPM statistics are in Supplementary Tables 4-5. Using the MirGeneDB miRNA set at FDR < 1% (unadjusted P-value < 2.34 × 10^-4^) we observed miR-QTLs for 224 of 470 tested miRNAs (Figure 2A). The distribution of the conditionally-independent miR-QTLs per eMiR is similar to what we observed with miRBase. Specifically, of the 224 eMiRs, 75% had one conditionally independent miR-QTL, 18% had 2, and about 6% had 3 or 4.

**Figure 2.**
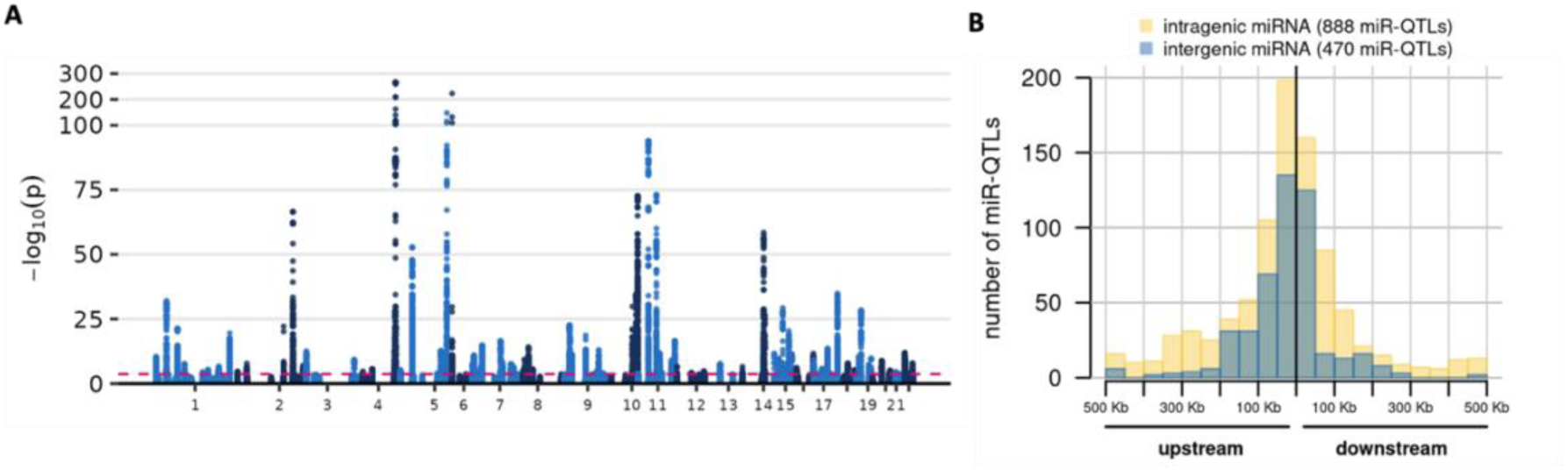
Summary of miR-QTL results. A) Manhattan plot of miR-QTL results. Each point represents a SNP-miRNA association. Pink dashed line indicates FDR of 1%. The Y-axis scale is condensed for -log_10_(P) ≥ 100. B) Distribution of miR-QTL position relative to the miRNA precursor for 1,358 index miR-QTLs. The miR-QTLs were stratified by whether the associated miRNA is intragenic (yellow) or intergenic (blue) and the two histograms are overlaid. The plot excludes 21 index miR-QTLs that had ambiguous relative position (because they were between or among precursors), were within the precursor, or were associated with miRNAs that could not be classified as intergenic or intragenic.

Notably, MirGeneDB miRNAs captured over 99.7% of the reads that mapped to the miRBase miRNAs, indicating that the miRNAs unique to miRBase are sparsely observed in our sequence data. Furthermore, the tested miRNAs in the MirGeneDB set included 75% (470 out of 624) of the tested miRNAs from the miRBase set. This substantial overlap in the tested miRNAs may be surprising at first given that miRBase lists several times more miRNAs than MirGeneDB. The simplest explanation is that miRBase-specific miRNAs are rarely detected in our data at or above our minimum required abundance threshold. Consistent with this, the tested miRNAs specific to miRBase tended to have lower expression than those present in both databases (median RPM of 3 versus 51, two-sided Wilcoxon rank-sum test P-value = 7.8 × 10^-31^).

The mapped read counts and miR-QTL summary statistics for both the miRBase and MirGeneDB miRNA sets are made available for researchers (see Data Availability section). For the downstream analyses in the rest of the manuscript, we opted to use the miR-QTLs from the MirGeneDB miRNAs to focus on miRNAs that all have the same high confidence support.

### 2.2 miR-QTL characteristics

#### 2.2.1 Percent expression variance explained

To recap, we observed miR-QTLs for 224 of 470 tested miRNAs from the MirGeneDB set (Figure 2A).

The median percent of miRNA expression variance explained by each conditionally independent miR-QTL was 4.1% (interquartile range 2.7%–7.8%), and 18 eMiRs have a miR-QTL that explains more than 20% of their expression variance. These values reflect not only the true effect size distribution of miR-QTLs but also the detection power in this study. We expect to have captured the strongest miR-QTLs; larger studies would be expected to identify additional miR-QTLs with smaller effect sizes.

#### 2.2.2 miR-QTL sharing by clustered miRNAs

Two unique features of miRNA biology can create correlation in the abundances of mature miRNAs. First, each miRNA precursor potentially encodes two mature miRNAs, from the -3p and -5p hairpin arms. Second, miRNA genes are often located in the genome in physical clusters and clustered genes may be transcribed as a unit. We can define a cluster as a group of miRNA genes separated by less than 10 Kb; some genes are ’standalone’ genes that are not located in a cluster. Using these definitions, the 224 eMiRs originate from 181 precursors that map to 135 distinct loci (either clusters or standalone genes). We observed moderate correlation between miRNAs from the same precursor or cluster. Specifically, the median pairwise Pearson correlation for miRNAs from the same precursor was 0.50 (interquartile range [IQR] 0.25- 0.68), and the median pairwise Pearson correlation for miRNAs in a cluster was 0.43 (IQR 0.27-0.70).

Predictably, miR-QTLs were sometimes, but not always, shared by miRNAs from the same precursor or cluster. For instance, 67% of SNPs tested for both the -3p and -5p miRNAs from the same precursor were associated with only one of the mature miRNAs.

#### 2.2.3 miR-QTL relative distance to eMiR genes

We characterized the position of the 1,384 index miR-QTLs relative to the respective precursor (or precursors if the miRNA has multiple precursors on the same chromosome as the miR-QTL). Three index miR-QTLs were located within the miRNA precursor sequence, including one located within the mature miRNA (Supplementary Table 7). About 9% of tested miRNAs mapped to more than one precursor; for one such miRNA, the two index miR-QTLs were located between precursors in a cluster, so we could not assign them to a single relative position category. Among the remaining miRNAs, miR-QTLs were more likely to be upstream than downstream of the miRNA gene (809 versus 570, two-sided binomial test P-value = 1.32×10^- 10^) (Figure 2B, Supplementary Table 8). As expected, miR-QTLs tended to be close to the gene’s transcription start site (TSS), and the distance to TSS was significantly correlated with P-value (Spearman rho 0.23, P- value = 1.14×10^-17^). We also observed that miR-QTLs for intragenic miRNAs tended to be further from the miRNA precursor than miR-QTLs for intergenic miRNAs (two-sided Wilcoxon rank-sum P-value = 6.31×10^-12^).

### 2.3 Extent of colocalization of miR-QTLs with host gene eQTLs and pQTLs

About 60% (or 135) of the eMiRs originate from intragenic miRNA precursors. Prior work has shown that intragenic miRNAs are sometimes co-expressed with their host genes^18,19^. Consistent with these reports, we found that the abundance of intragenic miRNAs was more likely to be positively correlated with abundance of host gene transcripts than expected by chance using matched dlPFC miRNA and transcriptomic data from 297 participants (two-sided Wilcoxon rank-sum P-value = 5.6×10^-13^). To follow up on this observation, we used coloc^20^, a Bayesian colocalization method, to identify specific instances of genetic co-regulation of intragenic miR-QTLs and their respective host genes. We combined the miR-QTL results with dlPFC eQTL observations from 621 ROS/MAP participants^21^. Considering the 106 miRNA-host pairs (103 mature miRNAs from 77 precursors in 71 host genes) for which the miRNA had miR-QTLs and the host gene had eQTLs, we found evidence of colocalization between miR-QTLs and host gene eQTLs (coloc PP.H4 > 0.5) for 17 pairs and no evidence of colocalization for 89 pairs (Supplementary Table 9). One interpretation of this finding is that in most cases, genetic regulation of miRNA expression and host gene expression occur independently. Alternatively, it could be that transcription is co-regulated, but post-transcriptional regulation of either the miRNA or host gene breaks the association between miRNA and host gene abundance. In any case, the findings show that for the majority of intragenic miRNAs, the miR-QTL signal is not explained by host gene eQTLs, and miR-QTLs provide information that is complementary to that from eQTLs.

Given the general tendency for positive correlation between miRNA and host gene transcript abundance, it is notable that two of the miRNA-host gene pairs with colocalized QTLs had significant negative correlation. For one of these pairs, miR-149-5p and host gene GPC1, previous work has shown that the miRNA targets the host gene in endothelial cells, has its own intronic promoter, and is activated independently of the host gene^22^. Our results together with the published findings suggest that the GPC1 eQTL signal may be driven by the miR-QTL effect on miR-149-5p and demonstrates that considering miR-QTLs can reveal the mechanism of action of host gene eQTLs. For the second pair, miR-1908-5p and host gene FADS1, we found no published experimental validation of the interaction. However, the Mendelian randomization analysis we describe in section 2.3 support this interaction.

In addition to testing colocalization between miR-QTLs and host gene eQTLs, we also had proteomics data available for 52 of the 106 miRNA-gene pairs to test colocalization between miR-QTLs and host gene pQTLs (Supplementary Table 10). Colocalization or lack thereof was consistent at the eQTL and pQTL level for 37 miRNA-host gene pairs, while colocalization was found at the eQTL level only for 8 pairs and pQTL level only for 5 pairs. Previous studies have found weak correlation between transcript and protein abundance^23–26^, so the discordance may be explained by multiple factors. For example, colocalization at the eQTL level only may be observed due to post-transcriptional regulation of either miRNA or host gene product.

### 2.4 Association of miR-QTLs with abundance of downstream target proteins

miRNAs are of interest because of their potential role in regulating abundance of their target genes. Although regulatory networks are complex and miRNAs are unlikely to be the sole regulatory mechanism for any given gene, we hypothesized that for at least a fraction of the eMiRs we could detect miRNA-QTLs that are associated with the protein expression of the eMiR’s target genes. In other words, we expected to find a small number of miR-QTLs that are also pQTLs (cis- or trans-pQTLs) for the miRNA’s target genes. To test this, we first obtained predicted target genes for each eMiR from TargetScan 7.2. We then tested whether the miR-QTLs for each eMiR were pQTLs for the eMiR’s predicted target genes using genotype and dlPFC proteomic data from up to 716 participants of ROS/MAP and the Arizona Study of Aging. These data included 511 donors that were also part of our miR-QTL dataset.

There were 206 eMiRs for which predicted downstream target genes were profiled in our dlPFC proteomics dataset. The median number of predicted targets per miRNA was 249 (interquartile range 165– 364). For all except 44 of ∼59,000 miRNA-protein pairs, the protein is trans (> 500 Kb from or on a different chromosome) to the miRNA. Therefore, for nearly all tests, we are effectively conducting trans-pQTL analysis. Recent work suggests that trans-QTLs collectively contribute substantially to heritability of complex traits, but each individual QTL is expected to have small effect^27^. The power to detect them is therefore limited at current sample sizes, especially when conducting naïve genome-wide searches due to the heavy multiple testing burden. By narrowing the pQTLs tests to SNPs for which we have some prior belief that they may be pQTLs based on the miR-QTL status and target prediction, we reduce the multiple testing burden.

Of the ∼59,000 miRNA-predicted target pairs (involving 206 miRNAs and 7,999 proteins), we discovered sharing of miR-QTLs and pQTLs for 11 pairs (Supplementary Table 11). These 11 pairs involved 10 mature miRNAs from nine precursors, and 10 unique target genes. One miRNA had pQTLs for two genes (miR-1307-5p with LYPLA2 and PPFIA3), and two miRNAs from the same precursor (miR-1307-3p and -5p) both had pQTLs for predicted target PPFIA3. For eight of the eleven pairs, the QTL SNPs are trans to the target gene. While the QTL sharing we detected is modest, this is not surprising given the expected small effect sizes for trans-pQTLs.

We further characterized the relationship among miR-QTL, miRNA expression, and protein expression for these 11 miRNA-target pairs using the SMR and HEIDI tests^28^. Like other Mendelian randomization-based methods, SMR aims to test the effect of an exposure (here, miRNA abundance) on an outcome (here, protein abundance) using SNPs as instrument variables. The SMR and HEIDI tests are combined to distinguish instances of causality (the SNP effect on protein expression is mediated by miRNA expression) and pleiotropy (the SNP directly affects both miRNA and protein expression) from linkage (distinct SNPs are each associated with miRNA or protein expression and the SNPs are in linkage disequilibrium). Five of the eleven miRNA-protein pairs had a significant SMR result after Bonferroni correction for the eleven SMR tests (P-value < 4.54×10^-3^), and four of these had HEIDI P-values that suggest the SMR signal is due to causality or pleiotropy rather than linkage (Table 1, Supplementary Table 12). For these pairs, we can consider the relative locations of the miRNA and target gene and the QTL effect directions to further interpret the relationship among the genetic variant and miRNA and protein abundances. Specifically, one pair comprises the intragenic miRNA miR-185-3p and its host gene, *TANGO2*. The top SMR SNP is located upstream of the host gene and has the same direction of effect on miRNA and protein abundance, so a plausible explanation is that genetic variation has a pleiotropic effect by affecting a regulatory element that co-regulates the miRNA and protein. The other three pairs comprise miRNAs and target genes located on separate chromosomes (Table 1). Given the low frequency of trans-acting regulatory regions, causality seems more likely than pleiotropy for these pairs. Furthermore, for these three pairs, the top SMR SNP is associated with miRNA and protein abundance in opposite directions, which is consistent with the QTL acting in trans on the protein abundance through its cis effect on miRNA abundance.

**Table 1.** miRNA and target gene pairs with evidence from SMR and HEIDI that genetically regulated miRNA abundance is associated with target protein abundance. RPM: reads per million.

| miRNA | miRNA median RPM | Target protein | miRNA position relative to target protein | miR-QTL effect direction | pQTL effect direction | SMR P-value | HEIDI P-value |
| --- | --- | --- | --- | --- | --- | --- | --- |
| miR-1307-5p | 48 | LYPLA2 | trans | down | up | 2.14E-10 | 0.64 |
| miR-1307-3p | 475 | PPFIA3 | trans | down | up | 9.25E-08 | 0.36 |
| miR-1307-5p | 48 | PPFIA3 | trans | down | up | 1.36E-07 | 0.42 |
| miR-185-3p | 34 | TANGO2 | intronic, same strand | up | up | 1.46E-07 | 0.39 |

The four miRNA-target pairs are not found in TarBase^29^, a database of experimentally validated miRNA-target gene interactions. To experimentally validate the potential interaction of the miRNAs and predicted target genes, we conducted luciferase assays in HEK 293T cells using the gene 3’ UTR reporter constructs and miRNA mimics. The results support interaction for all four pairs (Supplementary Figure 2), adding plausibility to the causal relationships inferred from the bioinformatic integration analyses.

TargetScan predicts targets only in the 3’ UTR, but targets can occur in other parts of the transcript. Moreover, target prediction tools use varying prediction criteria. Therefore, to expand on the above examination, we additionally collected miRNA-target interactions predicted by RNA22^30^ v2 and repeated the pQTL and SMR/HEIDI tests. The number of predicted interactions increased substantially, from ∼59,000 to ∼184,000, and we identified two additional miRNA-target pairs for which the SMR/HEIDI tests support pleiotropy or causality (Supplementary Table 12). In both cases, the top SMR SNP is associated with the miRNA and protein abundance in the same direction, which is not consistent with a causal role for the miRNA. In sum, the three trans miRNA-target pairs noted in Table 1 remain the strongest candidates for miRNAs that mediate the trans-pQTL effect on their predicted target gene.

### 2.5 miR-QTL enrichment in gene promoters and enhancers

We hypothesized that miR-QTLs were enriched in regulatory regions. To address this question, we intersected the miR-QTLs with cell-type specific promoters and enhancers identified for neurons, astrocytes, oligodendrocytes, and microglia by Nott et. al^31^.

We found that miR-QTLs were enriched (P-value < 0.05) in brain promoters combined across all four cell types (Figure 3). The Nott et al. study observed that promoters were largely shared across the four brain cell types^31^. Consistent with this observation, of the 1,272 miR-QTLs that overlap promoters, 61% overlap promoters in all four cell types. When we tested promoter enrichment for each cell type separately, the enrichment was not statistically significant for astrocyte and neurons, but a consistent trend was observed across cell types (Figure 3).

**Figure 3.**
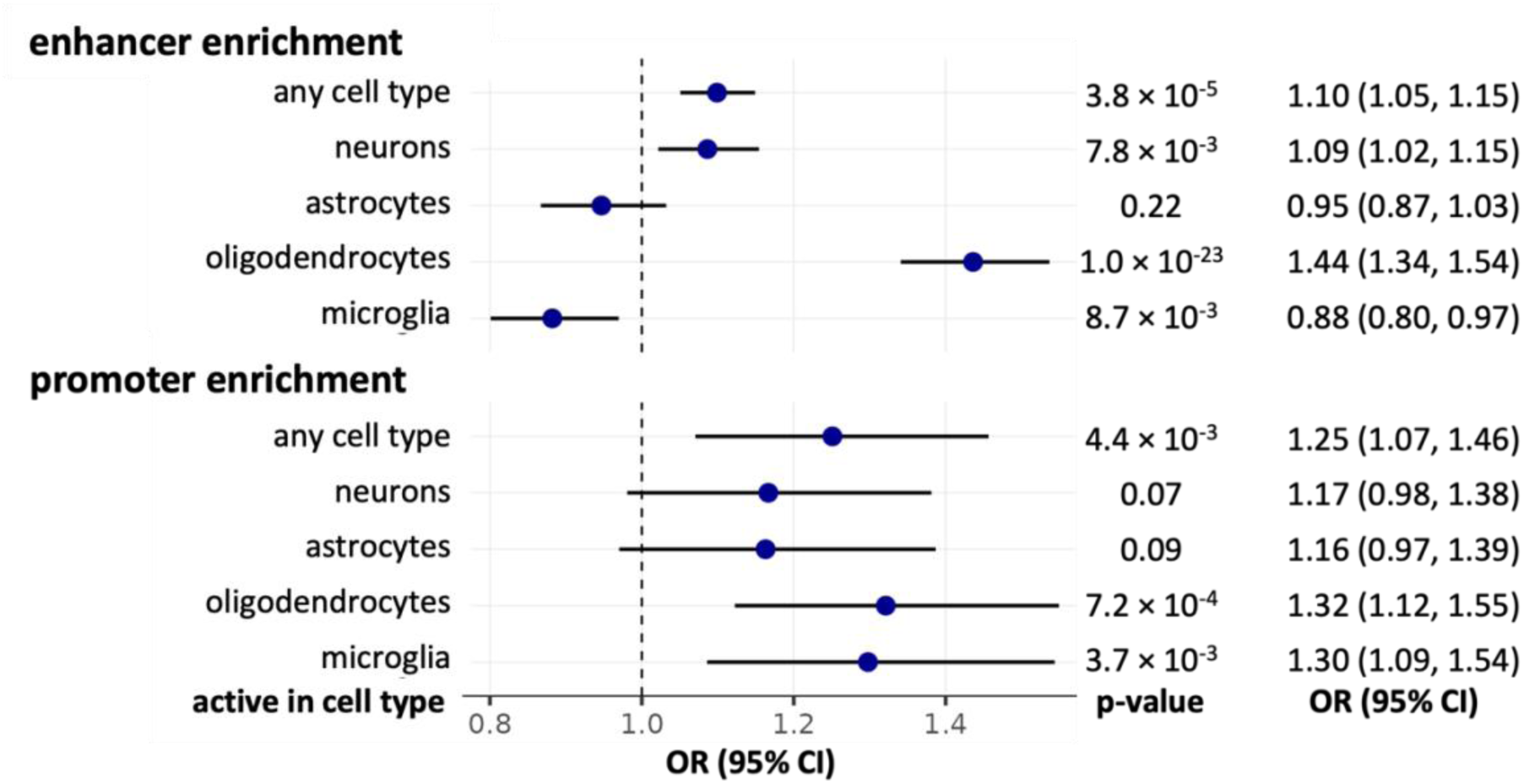
miR-QTL enrichment in brain enhancers and promoters. Enrichment was evaluated with the Mantel-Haenszel test. OR=odds ratio; CI=confidence interval.

We found that miR-QTLs were enriched in the union set of enhancers across all four cell types (Figure 3). In contrast with promoters, enhancer activity was observed to be largely cell type-specific. Considering enhancers for each cell type separately, miR-QTLs were enriched in enhancers in oligodendrocytes and neurons but not astrocytes; on the other hand, miR-QTLs were depleted in microglia enhancers (Figure 3).

We also tested miR-QTL enrichment in the reference enhancer list from PyschEncode, which was generated from bulk prefrontal cortex tissue^32^. miR-QTLs were enriched in this set of enhancers (Mantel-Haenszel common odds ratio = 1.32, 95% CI = [1.20,1.45], *p* = 1.3 × 10^-8^), and the odds ratio falls within the range of the odds ratios observed for the cell-type specific enhancer enrichments. We hypothesize that the microglia depletion may be an artifact caused by the fact that microglia comprise only a very small proportion of the cells used for our bulk miR-QTL analysis. Consequently, genetic variants that affect miRNA abundance in microglia only (e.g., because they affect activity of microglia-specific enhancers) will have lower detection rate than miR-QTLs with signal coming from any of the cell types that are more prevalent in our sample. The combination of this bias with the cell-type specific nature of enhancers could manifest as lower miR-QTL rate in microglia enhancers compared to non-microglia enhancers.

### 2.6 Cross-tissue comparison of miR-QTLs

We compared our miR-QTL results to those from two recent studies performed in other human tissues. One study used qRT-PCR to measure miRNA expression in whole blood from up to 5,239 (minimum 200) participants of the Framingham Heart Study and identified cis-miR-QTLs (within 1 Mb of the mature miRNA) for 76 of 280 miRNAs^33^. Another study analyzed small RNA-sequencing from 212 fetal brain neocortex samples and identified cis-miR-QTLs (within 1 Mb of the miRNA precursor) for 65 of 811 mature miRNAs at their stringent threshold^34^.

To understand the rate of eMiR sharing between our study and each of the other two studies, we compared the eMiR counts accounting for the miRNAs that were tested in each study (Supplementary Figure 3). We first compared our eMiRs to those discovered in blood. Of the 224 eMiRs we identified, 73 were tested in whole blood and 41% of those (30 of 73) were identified as eMiRs in that context. Conversely, of the 76 eMiRs identified in whole blood, 55 were tested in our study and we identified 55% of those (30 of 55) as eMiRs (Supplementary Figure 3). A chi-squared test using the 173 miRNAs tested in both studies indicates that the number of shared eMiRs is higher than expected by chance, *X*^2^(1, *N* = 173) = 4.3, *p* = .038. It is important to recognize that the detected miR-QTLs in each study reflect not only differences in miRNA genetic regulation across contexts but also the sensitivity and specificity of each study. Performing the same comparison between our results and the fetal neocortex results, we found that only 13% (23 of 184) of the eMiRs we identified were also identified as eMiRs in the fetal neocortex study. Yet, most fetal eMiRs were evident in our study (74% [23 of 31]) (Supplementary Figure 3). These results suggest differences between the studies likely stem from differences in sample sizes. A chi-squared test using the 394 miRNAs tested in both studies indicates that a miRNA that is an emiR in one study is more likely to be an emiR in the other study, *X*^2^(1, *N* = 394) = 9.1, *p* = 2.6 × 10^-3^. Taken together, we observe significant sharing of miRNA genetic regulation between contexts, but important differences in study design and power limits further inference.

### 2.7 Uncovering miRNAs contributing to the pathogenesis of brain disorders

GWAS have identified many risk loci for brain illnesses; however, how these loci confer risk in these psychiatric and neurologic diseases remain unclear. Recently, integrating GWAS signals with gene expression and QTL data from a reference set has become a powerful strategy for identifying genes that are associated with complex traits in the absence of expression measures in the larger GWAS participants. Here, we leveraged our miRNA expression data and miR-QTL results to uncover genetic variants that confer risk for psychiatric or neurodegenerative diseases through their effects on brain miRNAs.

To do so, we integrated the genome-wide miRNA profiles with results from GWAS of these brain traits using multiple independent and complementary approaches (Figure 1). First, we performed a miRNA- wide association study (miRWAS) using FUSION^35^, utilizing the miR-QTL results from the reference ROS/MAP dataset to impute genetically regulated miRNAs in the larger GWAS participants and then identify miRNAs that are associated with the brain trait. Next, we determined whether the identified miRNAs and the brain trait share causal variants through colocalization analysis using coloc^20^. Subsequently, we performed SMR^28^ to verify that the identified miRNAs mediated the association between the GWAS loci and the brain trait and ruled out association due to linkage disequilibrium with the HEIDI test^28^ (Figure 1). In other words, a miRNA was declared as being consistent with a causal role in a brain trait if it met all of the following four criteria: i) the miRNA was associated with the brain trait in the miRWAS at FDR P-value < 0.05; ii) the identified miRNA and brain trait share a causal variant as reflected by the coloc posterior probability of a shared causal variant (PP.H4 >0.5); iii) the miRNA showed evidence for mediating the association between the GWAS loci and the brain trait in the Mendelian randomization analysis, and iv) the mediation was not due to linkage disequilibrium based on the HEIDI test^28^. These miRNAs are referred to as causal miRNAs for simplicity henceforth.

We used data from the latest GWAS summary statistics in five neurodegenerative diseases (Alzheimer’s disease^36^, frontotemporal dementia^37^, amyotrophic lateral sclerosis^38^, Lewy body dementia^39^, and Parkinson’s disease^40^) and eleven psychiatric disorders (major depressive disorder^41^, bipolar disorder^42^, schizophrenia^43^, anxiety^44^, post-traumatic stress disorder^45^, alcoholism^46^, neuroticism^47^, insomnia^48^, attention deficit hyperactivity disorder^49^, autism^50^, and suicide attempt^51^; Supplementary Table 13). Among the 470 available miRNAs, 140 had significant heritability (defined as heritability p < 0.01) and were included in the miRWAS (Supplementary Table 14).

We identified 49 unique miRNAs associated with one or more brain traits in the miRWAS (Figure 4; Supplementary Tables 15-16). Further imposing the remaining three criteria described above, we found four unique miRNAs to be consistent with a causal role in Parkinson’s disease (miR-92b-3p), bipolar disorder (miR-1908-5p and miR-499a-5p), major depressive disorder (miR-1908-5p), post-traumatic stress disorder (miR-190b-5p), and schizophrenia (miR-499a-5p) (Table 2). Notably, two miRNAs were causal in two brain traits—miR-1908-5p in both bipolar disorder and major depressive disorder and miR-499a-5p in both bipolar disorder and schizophrenia (Table 2).

**Figure 4.**
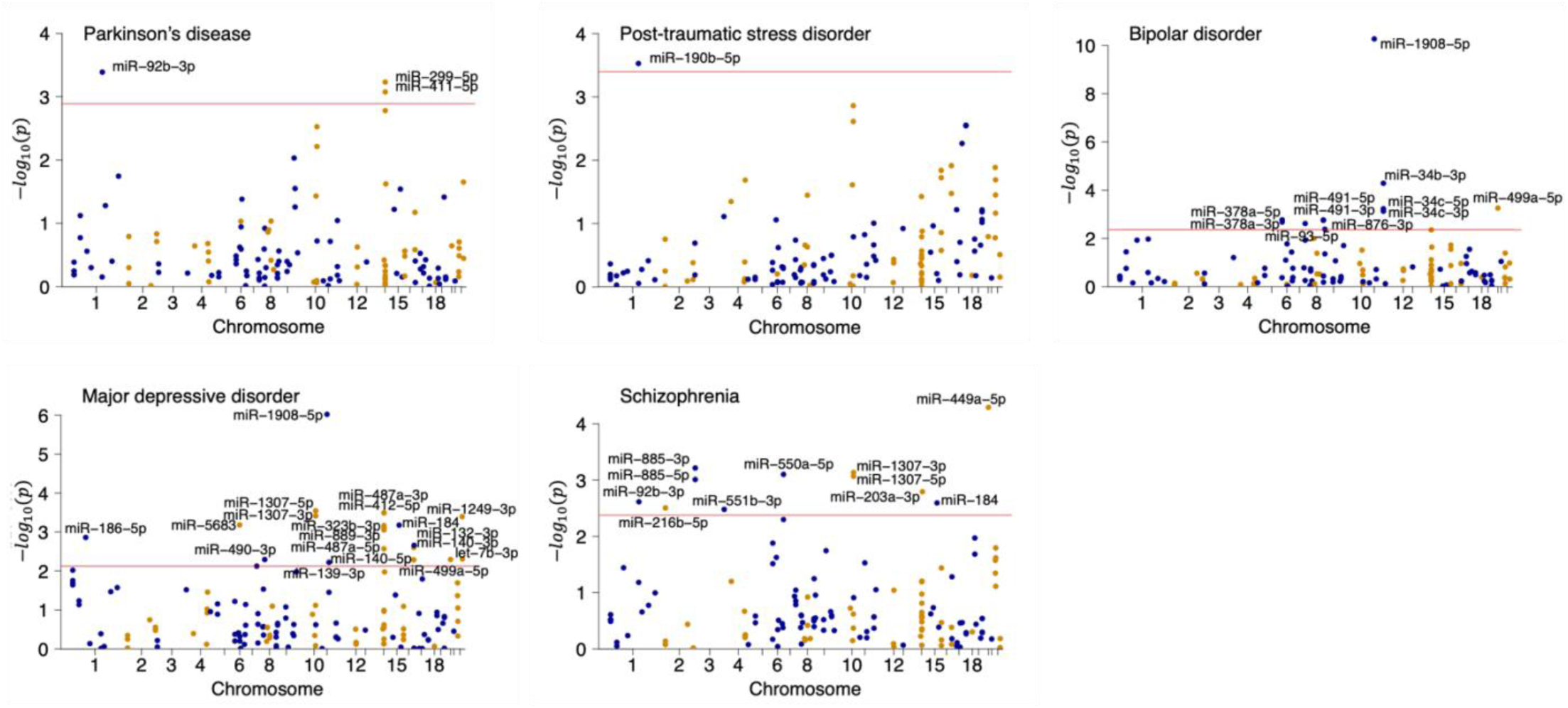
Manhattan plots for the five traits with associated miRNAs in the miRWAS analysis. Associated miRNAs (at false discovery rate < 5%, indicated by the horizontal red line) are labeled.

**Table 2.**
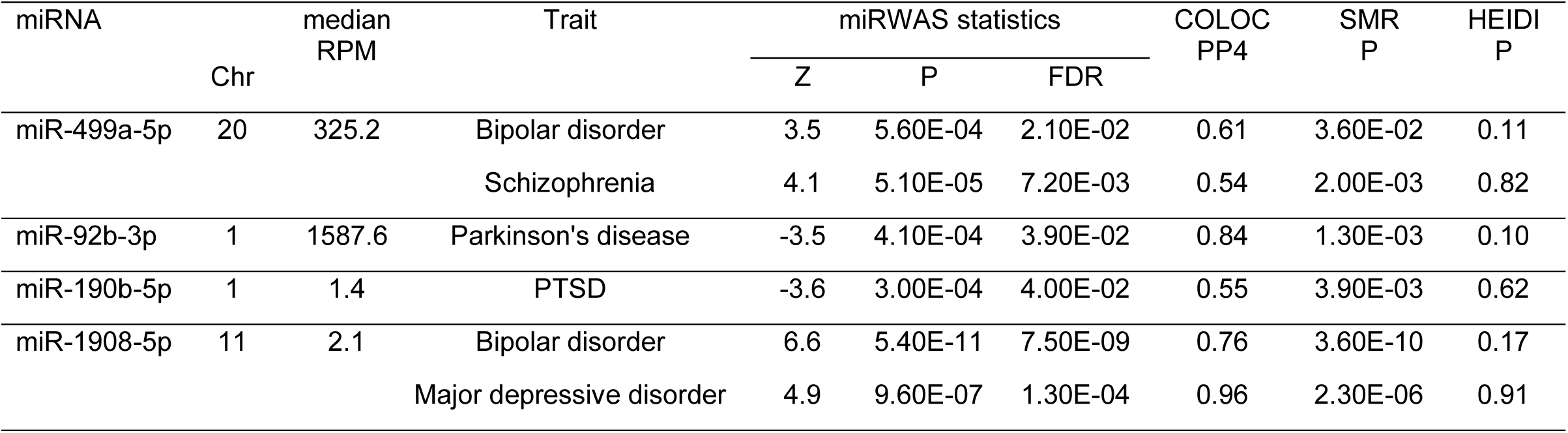
Causal miRNAs in psychiatric and neurodegenerative diseases. . PTSD: post-traumatic stress disorder.

### 2.8 Examining multi-omics signals in brain disorders

Since miRNAs may share QTLs with nearby genes due to linkage disequilibrium, pleiotropy, or mediation, we examined whether the causal role could be attributed to the above identified miRNAs instead of their nearby genes. We identified a set of causal transcripts and proteins for a range of psychiatric and neurodegenerative diseases in a recently published work^26^. In that work, we integrated GWAS summary statistics for the brain traits with brain transcriptomes (n=888) and proteomes (n=722) using TWAS/PWAS, Mendelian randomization, and colocalization analyses to identify mRNAs and proteins consistent with a causal role in the brain traits^26^. Inspecting the four causal miRNAs we identified here, we found that three causal miRNAs are located in proximity (< 500Kb) to one or more of the causal transcripts or proteins identified in the prior work.

Specifically, miR-1908-5p, causal in both bipolar disorder and major depression, is located within FADS1 and is close to TMEM258, which are causal mRNAs in both bipolar disorder and major depression^26^ (Figure 5; Supplementary Table 17). Using SMR for two molecular traits, we found evidence that miR-1908- 5p mediates the association between its miR-QTLs and FADS1 but does not mediate the association between its miR-QTLs and TMEM258 (Supplementary Table 17). As mentioned in section 2.2, miR-1908-5p miR-QTLs colocalize with FADS1 eQTLs and the miRNA and transcript abundances are negatively correlated. These findings suggest that miR-1908-5p likely contributes to bipolar disorder and major depression through its effect on FADS1 mRNA expression but acts independently from TMEM258. Analogously, we found that miR- 499a-5p is an independent causal mediator from GGT7 and EDEM2 in bipolar disorder (Supplementary Figures 4-5; Supplementary Table 17). Likewise, miR-92b-3p is an independent casual mediator from EFNA3 in Parkinson’s disease (Supplementary Figure 6, Supplementary Table 17). In summary, except for miR-1908- 5p, the other three causal miRNAs are causal mediators independent of nearby causal transcripts and proteins and provide additional valuable biological insights into the pathogenesis of these psychiatric and neurodegenerative diseases.

**Figure 5.**
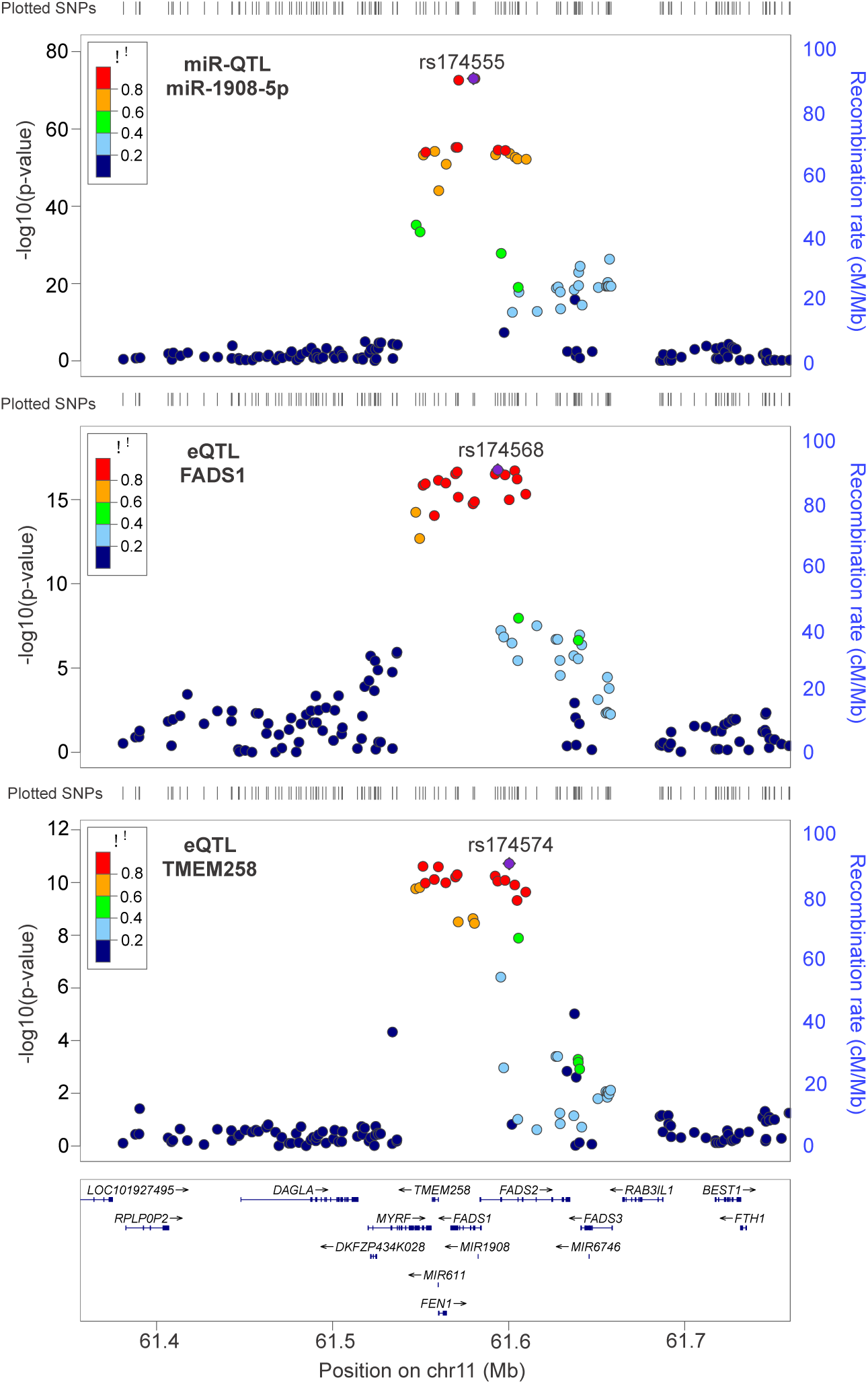
LocusZoom plots for miR-1908-5p miR-QTLs and FADS1 and TMEM258 eQTLs. In each plot, the SNP with the lowest P-value is colored purple, and the remaining SNPs are colored according to the extent of LD with that SNP. SMR and HEIDI analysis found that miR-1908 miR-QTLs colocalized with FADS1 eQTLs, but not TMEM258 eQTLs.

## 3 Discussion

Using more than 600 small RNA-sequencing profiles from dlPFC of older adults, we identified miR-QTLs for 224 miRNAs and integrated those results with human brain proteomes and GWAS summary statistics to investigate how genetically regulated brain miRNAs impact protein expression and neurodegenerative and psychiatric traits. While genome-wide miR-QTL mapping efforts have been conducted in various human tissues (blood^52,53^, fetal brain^34^, adipose^54^, liver^55^), to our knowledge the results we report here are the first set of miR-QTLs available for adult human brain.

Importantly, we performed parallel runs of the miR-QTL analysis using reads mapped independently to two sets of miRNA annotations – either the popular miRBase database (Release 22.1) or the curated MirGeneDB database (version 2.1). MirGeneDB contains less than 30% of the annotations in miRBase due to its more stringent inclusion criteria. However, since we filtered both annotation sets using a minimum abundance threshold and since annotations specific to miRBase tended to be lowly expressed in our brain samples, there was a much larger overlap in the tested miRNAs, with 75% of miRNAs tested in both runs. We chose to use the MirGeneDB set for the subsequent colocalization and integration analyses since including poor quality annotations would reduce the power and interpretability of these analyses; however, we recognize that the more comprehensive miRNA set may be valuable in other contexts and make all our data available as a resource for future studies.

We identified cis-miR-QTLs for almost half of the miRNAs that were tested, which is comparable to 53% of the proteins having pQTLs at FDR < 1% in a recent published study in 716 individuals^26^. Since many miRNAs are located within host genes, a question that arises is how often these intragenic miRNAs are co-regulated with their host genes. This question has been investigated previously using various approaches such as examining TSS sharing and evaluating co-expression in cell lines^56^. Our colocalization analysis, which utilized population variation in miRNA expression, host gene expression, and genotypes, complements other approaches. Notably, we found that the majority of intragenic miRNAs were not genetically co-regulated with their host genes, which may indicate that miRNAs have biological roles distinct from the functions of their host genes. It also demonstrates that even for miRNAs located within protein-coding genes, miR-QTL mapping detects potentially important genetic variants that would not be found in other QTL studies.

QTLs for master regulators such as miRNAs can potentially have an outsized effect since they can influence the expression of many other genes. We leveraged our miR-QTL results to identify brain traits and proteins that are influenced by genetically regulated miRNA abundance. We identified four miRNAs consistent with a causal role for complex brain traits. Our findings that miR-1908-5p and miR-499a-5p are causal in bipolar disorder are consistent with those from a prior study that used a candidate gene-based association study approach^57^. Notably, the miR-92b we identified to be causal in Parkinson’s disease has been shown to target PTEN, promote neurite development^58^, and maintain neuroblast self-renewal^59^. Our study complements this existing literature and is the first to use human brain miRNA expression profiles to provide support for a causal role of these miRNAs. We present here the first evidence for the contribution of miR-1908-5p in major depression, miR-190b-5p in post-traumatic stress disorder, miR-499a-5p in schizophrenia, and miR-92b-3p in Parkinson’s disease, to our knowledge. These causal miRNAs present new avenues to elucidate the molecular mechanisms underlying these neurodegenerative and psychiatric conditions.

Considering pairs of miRNAs and predicted target genes, we observed 18 miRNA-target gene pairs (involving 16 unique miRNAs) for which the miR-QTLs were also pQTLs for the predicted target in the same tissue. Using the SMR and HEIDI tests, we found that six miRNA-target pairs were consistent with pleiotropy or causality. Our subsequent validation experiments confirmed potential interaction for all four pairs that were tested. While further investigation is necessary to confirm endogenous repression activity and the observed QTL associations, these findings are an initial demonstration of how miR-QTLs can be used to prioritize putative trans-pQTLs and provide additional support for predicted interactions.

We observed significant enrichment of miR-QTLs in promoters active in human brain^31^. We limited this analysis to SNPs within 50 Kb of miRNA genes because doing so reduces the potential effect of confounding between distance to nearest gene and probability that a genomic region harbors a promoter ^60^. We note that we searched only for direct overlap with promoters, and in a future analysis the miR-QTLs could be integrated with chromatin interaction maps to identify miR-QTLs that are distal to the promoter but are in regions that physically interact with it^61^.

Unexpectedly, we observed significant depletion of miR-QTLs in microglia enhancers. The promoter and enhancer detection was done using pooled sorted single nuclei^31^, so the relative proportions of the cell type in bulk tissue does not create a bias in the promoter and enhancer distributions across cell types. Our analysis, on the other hand, used bulk tissue, and microglia are a minor cell type in our samples. The observed depletion of miR-QTLs in microglia enhancers is likely an artifact arising from the combination of the enhancers being largely mutually exclusive across cell types, the low power to detect miR-QTLs specific to microglia cells, and the enrichment of miR-QTLs in enhancers of other cell types. Also, the promoter and enhancer annotations were made using data from only 10 individuals and may have missed regulatory regions whose activity varies across individuals^31^.

Comparing our miR-QTLs with miR-QTLs detected in fetal (midgestation) neocortex and adult whole blood showed higher eMiR sharing than expected by chance. However, the three studies have important differences, such as sample size, approach to miRNA quantification, number of miRNAs considered, and analytic approach, that influence miRNA-QTL detection. Differences in power due to sample size are likely the most salient and make direct comparisons challenging to interpret. Large samples from across the age and tissue spectrum will help clarify the eMiRs that are unique or common and elucidate the context specificity of miR-QTLs.

We recognize several limitations of this study. First, the findings would be strengthened by replication in an independent sample. The power of this analysis is tied to the relatively large sample size for the miRNA profiles, and there is not an independent comparable brain miRNA dataset for replication at this time.

Second, although we characterized miRNA expression in a relatively large sample, all the participants were of European ancestry, and expanding studies to include people of diverse ancestry is imperative. Third, brain miRNA genetic regulation may differ across stages of adulthood and ought to be tested in future studies that include participants from a wider age range. Nevertheless, our observations fill a gap in the available resources for studying the role of miRNAs in the human brain. Fourth, our analyses were conducted using samples from a specific brain region (dlPFC) and our observations may not be characteristic of other brain regions. Fifth, some of the tested miRNAs had modest expression in our dlPFC bulk tissue samples and their cell type-specific expression levels need to be quantified in relevant cell types. Notable among those, miR- 1908-5p was previously observed to be differentially expressed in blood in the acute bipolar state compared to remission state^62^, which suggests that miR-1908-5p is relevant to bipolar disorder despite its relatively low abundance. Finally, since we focused the analyses on the manually curated miRNAs present in MirGeneDB, we may have missed some true miRNAs that did not meet the criteria of that database. The trade-off for this choice is that we have high confidence in the validity of the miRNAs we did consider, which improves the signal in our observations and increases the confidence that our results provide a solid foundation to build on in future downstream analyses^11^.

We also highlight some of the strengths of this study. First, we minimized the impact of cell-type variation by estimating surrogate variables and including these as covariates in the regression models for miR-QTL mapping. These surrogate variables were designed to capture as much as possible the potentially unknown, unmeasured, or hidden confounding variables, including cell type heterogeneity. Second, we examined the relationship of genetic regulation of miRNAs and their downstream targets using proteomic data. Proteomic data may better capture the effect of miRNA post-transcriptional regulation on target genes than transcriptomic data, which is more commonly used. Finally, this is the largest set of genome-wide miRNA sequencing profiles from human brain to the best of our knowledge. This enabled us to conduct well-powered miR-QTL analysis and to successfully integrate these results with GWAS data to identify miRNAs contributing to the pathogenesis of psychiatric and neurodegenerative diseases. The miRNA expression data as well as the full catalog of miR-QTL results will be available to the research community for further studies. Given the role of miRNAs as critical regulators, these data represent a valuable resource for studies aimed at characterizing the complex networks that link population genetic variation to brain-related traits and disease.

## 4 Online Methods

### 4.1 ROS/MAP cohort and phenotype data

The Religious Orders Study (ROS) and Rush Memory and Aging Project (MAP) are U.S.-based longitudinal community-based cohort studies of cognitive decline and aging^63^. ROS enrolls priests, monks, and nuns from sites across the country, while MAP enrolls individuals from retirement communities, social service agencies, and church groups in the greater Chicago area. All participants provided informed consent, underwent annual cognitive evaluations, were organ donors, and signed an Anatomical Gift Act and repository consent to allow their data and biospecimens to be repurposed. An Institutional Review Board of Rush University Medical Center approved both studies and we have complied with all relevant ethical regulations. Clinical diagnosis at death was based on the opinion of a neurologist with expertise in dementia who had access to select clinical but not postmortem data. Sex (male or female) was based on self-report. Post-mortem interval (PMI) is the time interval from time of death to autopsy. Self-identified race was defined based on the participant’s response to the question ‘What is your race?’ With possible answers White; Black or African American; American Indian or Alaska Native; Native Hawaiian or Other Pacific Islander; Asian; or Other.

### 4.2 MiRNA data

#### 4.2.1 RNA extraction from frozen brain tissues

Frozen post-mortem dorsolateral prefrontal cortex tissue samples from 743 brains were obtained from Rush University. RNA extraction from the tissue samples was performed with TRIzol (Invitrogen, Carlsbad, CA, USA) following the manufacturer’s protocol. Approximately 80mg (ranging 50mg to 100mg) of frozen brain tissues were used to extract RNA.

#### 4.2.2 RNA quality control and measurement

RNA quality was measured by two methods following the manufacturer’s protocol: 1) Nanodrop Spectrophotometer (Thermo Fisher Scientific, Waltham, MA, USA) 2) Agilent Bioanalyzer RNA 6000 Nano (Agilent Technologies, Santa Clara, CA, USA). Nanodrop Spectrophotometer measures concentration and A_260_/A_280_ ratio which determines the purity of RNA. The average concentration of extracted RNA samples was around 900ng/µl resuspended in 50µl DEPC-treated water, and the average A_260_/A_280_ ratio was 1.93. Agilent Bioanalyzer RNA 6000 Nano was used to determine an RNA Integrity Number (RIN) to assess quality RNA. The average RIN score for all 743 RNA samples was 5.5; 672 samples with RIN ≥ 2.1 were selected for miRNA library preparation. Based on RIN, PMI, sex, cognitive diagnosis, amyloid, tangles, depression, education, and sense of life purpose, samples were randomized into batches of 8 for miRNA library preparation.

#### 4.2.3 miRNA library preparation

miRNA library preparation was conducted using New England Biolabs’ (NEB) NEBNext Muliplex Small RNA Library Prep Kit for Illumina Index Primers 1-48 (NEB, Ipswich, MA, USA) following the manufacturer’s protocol. 1 µg of total RNA was used as input for the miRNA library preparation in batches of 48. Libraries were checked for the presence of a 21-nucleotide insert using an Agilent Bioanalyzer DNA 1000 prior to size selection with a Pippin Prep (Sage Science, Beverly, MA, USA).

#### 4.2.4 miRNA library pooling and sequencing

The molarity of each miRNA library was measured by Qubit Fluorometer (Thermo Fisher), and libraries were diluted and pooled. MiRNA sequencing was performed using an Illumina HiSeq 3000 (Illumina, San Diego, CA, USA) at the Emory Yerkes Genomics Core (Atlanta, GA).

#### 4.2.5 miRNA quantification, normalization, and quality control

Adaptor sequences were trimmed using Trimmomatic^64^ (version 0.36), and miRNA counts were generated using mirDeep2^65^. Specifically, we first used the “mapper.pl” from mirDeep2 to generate collapsed fasta files, then removed reads with length less than 20 nucleotides or greater than 25 nucleotides using a custom script. Then “quantifier.pl” from mirDeep2 was used to map the reads to known human precursor and mature miRNA sequences obtained from MiRBase^12^ (Release 22.1). Reads were allowed to map up to 2 nucleotides upstream of the mature sequence and up to 5 nucleotides downstream of the mature sequence, and with up to one mismatch. These are the default parameters for mirDeep2, and are reasonable because they allow for untemplated nucleotide addition and imprecise processing, which are common especially at the 3’ end of mature miRNAs. We removed very lowly-expressed miRNAs, which we defined as those with < 1 RPM for > 50% of samples. We chose a liberal threshold for this filter to allow for the fact that miRNAs can have low observed RPM in our heterogeneous bulk tissue samples but higher expression in specific cell types. For miRNAs mapped to multiple precursors, we kept the entry with the highest total count across samples.

Since miRBase has been reported to contain a substantial number of false positive miRNAs^66^, we performed a second mapping run after filtering the reference precursor and mature miRNA fasta files to only the 504 precursors and 857 mature miRNAs present in MirGeneDB^13^ (v2.1), a database of miRNAs that meet a set of criteria designed to distinguish true miRNA sequences from false annotations and other types of small RNAs^14^.

Sample filtering was performed on several criteria. First, we iteratively removed samples that were greater than 5 standard deviations (SD) from the mean within the respective batch for total read count, trimmed read rate, and mapped read rate. We then filtered 11 samples to focus analysis on individuals of European ancestry based on genetic data. The analysis was limited to these participants because of small sample sizes for participants from other genetic ancestries. The raw counts were then normalized for library size and transformed to log2 counts using EdgeR. Principal components (PCs) were estimated, and samples that were greater than 4 SD from the mean of either of the first two PCs were removed.

Surrogate variables (SVs) were estimated from the normalized counts using R package ‘sva’. The number of SVs to estimate was set at 15 based on the results of the num.sv function, and SVs were estimated with a randomly-generated binary variable as the variable of interest and miRNA batch, PMI, RIN, age at death, cognitive diagnosis at death, study (ROS v. MAP), and sex as adjustment variables.

### 4.3 Genotype data for miR-QTL calling and miRWAS

DNA was extracted from blood or brain and genotyping was performed using whole-genome sequencing or genome-wide genotyping (Affymetrix GeneChip 6.0 or Illumina OmniQuad Express platforms) as described previously^67^. Initial filtering was performed on the WGS, Affymetrix chip, and Illumina chip datasets separately^68^. Specifically, samples were excluded if they had genotype missingness >5% or were from participants who self-identified their race as other than White, and variants were excluded if they had evidence of deviation from Hardy Weinberg equilibrium (P-value < 1×10^-8^), genotype missingness >5%, minor allele frequency <1%, or were not a single nucleotide polymorphism (SNP). Sample outliers with respect to population structure were removed using EIGENSTRAT^69^. For samples with array-based genotypes, imputation was performed to 1000 Genome Project Phase 3^70^ using the Michigan Imputation Server^71^, and SNPs with *R*^2^>0.3 were retained.

Genotypes for the participants with miRNA data were then extracted from the WGS and imputed chip datasets and merged. Some participants were genotyped using both WGS and chip; in these cases the WGS genotypes were kept. Before merging, variants with MAF < 5% in each dataset were removed. We then used bcftools to identify sites common to both datasets and with identical reference and alternate alleles and merge the sample data for these sites. The minor allele frequency, Hardy-Weinberg equilibrium, and genotype missingness checks were repeated after merging. Second degree and closer relatives were removed using KING^72^. Sample characteristics for the 604 participants that were included in the analysis are summarized in Supplementary Table 1. We calculated the top ten genetic PCs using PLINK2^73^ to be used as covariates in the miR-QTL analysis. The PC analysis showed that PC1 separates samples based on genotype source (WGS v. array). PC analysis also confirmed that all samples cluster with the 1KG CEU population. For the miRWAS analysis, we restricted the SNPs to the set of 1,190,321 SNPs in the linkage disequilibrium reference panel provided by FUSION^35^.

### 4.4 Genotype and proteomic data for pQTL and SMR analysis

The pQTL analysis was conducted using SNP genotypes and dlPFC proteomic profiles from participants of the ROS/MAP studies and the Arizona Study of Aging and Neurodegenerative Disorders, conducted by the Banner Sun Health Research Institute. The Banner study primarily recruits cognitively normal individuals from Phoenix, AZ retirement communities, and also some individuals with AD or Parkinson’s disease^74^. Participants or their legal representatives sign an Institutional Review Board-approved consent form allowing for brain donation, use of donated biospecimens for approved future research, and genetic studies.

The ROS/MAP and Banner data were combined for this analysis. The data generation has been described previously (ROS/MAP proteomics^75^; Banner proteomics^76^, ROS/MAP genotypes^67^; Banner genotypes^76^). The quality control, normalization and data aggregation procedure have also been described previously^26^. Proteomic profiles were represented by residuals after regressing out effects of batch, PMI, age at death, and cognitive diagnosis. Of the 716 participants included in the pQTL and SMR analysis, 565 are from the ROS/MAP cohort and 511 of those overlap the 604 participants included in the miR-QTL analysis. Sample size varied across different proteins from 76 to 716 participants due to differences in missingness rate across proteins.

### 4.5 GWAS summary statistics

We used association summary statistics from GWAS studies of five neurodegenerative diseases (Alzheimer’s disease^36^, frontotemporal dementia^37^, amyotrophic lateral sclerosis^38^, Lewy body dementia^39^, Parkinson’s disease^40^) and eleven psychiatric traits (major depressive disorder^41^, bipolar disorder^42^, schizophrenia^43^, anxiety^44^, post-traumatic stress disorder^45^, alcoholism^46^, neuroticism^47^, insomnia^48^, attention deficit hyperactivity disorder^49^, autism^50^, suicide^51^). We focused on GWAS summary statistics results from samples of individuals with European ancestry (Supplementary Table 13).

### 4.6 Validation of miRNA-target interaction

To validate the predicted miRNA/mRNA target pair interactions, we transfected human HEK-293T cells with pairs of a miRNA mimic and a 3’-UTR reporter construct with firefly luciferase as a reporter, as follows: miR- 1307-5p with 3’-UTR of PPFIA3; miR-1307-3p with 3’-UTR of PPFIA3; miR-185-3p with 3’-UTR of TANGO2; and miR-1307-5p with 3’-UTR of LYPLA2. Catalog numbers for materials used in transfection and luciferase reporter assays are available in Supplementary Table 18.

Human HEK293T cells were maintained in DMEM (Dulbecco’s modified Eagle’s Medium), supplemented with 10% FBS and 1% antibiotics solution (Pen/Strep), at 37 °C and 5% CO2. One day prior to transfection, HEK293 cells were plated on two 96-well plates (50,000 cells/well). The cells were transfected using Lipofectamine 2000 (Thermofisher, MA, USA) with the miRNA mimic and a 3’-UTR reporter construct pairs as indicated above and incubated for 48 h at 37 °C and 5% CO2. Each miRNA mimic/3’-UTR reporter construct pair was transfected in quadruplicate on each plate. Luciferase assay was subsequently carried out with the ONE-Glo™ EX Luciferase Assay (Promega, WI, USA). Transfected cells were treated in accordance with the manufacturer’s protocols and luminescence was assayed on a Synergy HTX multi-mode microplate reader (Biotek, Winooski, VT, USA).

### 4.7 Statistical analysis

All statistical analyses were done using R version 3.6.0 unless otherwise noted. All genome coordinates are with respect to human reference GRCh37. Linear regression was performed to compare luminescence between experimental conditions adjusting for plate effects. Significant difference was defined as P-value < 0.05.

#### 4.7.1 miR-QTL mapping

MiRNA coordinates were obtained from miRBase v.22 and then converted from GRCh38 to GRCh37 using LiftOver to match the build used for the genotype data and GWAS summary statistics. MiRNAs included in the analysis met the abundance threshold described above, had gene coordinate information in the coordinate file, had at least one precursor gene located on an autosome, and had at least one common (MAF >5%) SNP within the test window. The test window for each mature miRNA included the precursor sequence(s) plus 500 Kb symmetric padding around each precursor sequence. For miRNAs with multiple precursors that were less than 500 Kb apart, SNPs that fell in the padding region for multiple precursors were tested only once for the miRNA.

Samples included in the analysis passed all genotyping and miRNA data QC, had normal cognition, MCI, AD, or other dementia at death, and were of European ancestry. The effect of SNP allele dosage (assuming an additive effect) on miRNA abundance was estimated for each SNP-miRNA pair in turn using the --glm function from PLINK2. The following variables were included as covariates: miRNA batch, PMI, RIN, age at death, cognitive diagnosis at death (normal v. impaired), study (ROS vs. MAP), sex, genotype platform, 10 genetic PCs, and 15 miRNA SVs. We observed that the genetic PCs are strongly correlated with genotype platform. We set the max variance inflation factor to 500 to allow the tests to run despite multicollinearity between these variables. Q-values were calculated using R package ‘qvalue’ using p-values from all tests. We define miR-QTLs as SNP-miRNA pairs with q-value < 0.01.

Since we performed the miR-QTL mapping using dense genotyping, many miR-QTLs are in high linkage disequilibrium (LD) and represent essentially the same genetic signal. We extracted a reduced set of miR-QTLs for each miRNA using the ‘--clump’ function from PLINK 1.9. This function applies a greedy algorithm to iteratively prune miR-QTLs to arrive at a set of ‘index’ miR-QTLs that have pairwise r^2^ < 0.5 or are separated by at least 250 Kb.

The percent of miRNA variance explained by individual miR-QTLs was estimated using R package ‘variancePartition’. The outcome in the model was the residuals after regressing the effects of miRNA batch, PMI, RIN, age at death, cognitive diagnosis, study, sex, and miRNA SVs from the normalized expression profiles, and the predictors were SNP and 10 genetic PCs.

For each miRNA with more than one index SNP, we used conditional analysis to identify miR-QTLs with independent signal. First, we ranked the miR-QTLs for each miRNA by increasing P-value. The independent variants list was initiated with the miR-QTL with the lowest P-value. For each remaining miR-QTL, an association test was run using the same model as the initial analysis except with the independent variant(s) also included as predictors. If more than one miR-QTL remained significant, the top ranked miR-QTL was added to the independent variants list and another round of testing was performed with the remaining miR-QTLs. If only one miR-QTL was significant, it was added to the independent variants list and the procedure was complete.

#### 4.7.2 MiRNA cluster and intra-/intergenic status

MiRNAs were defined as intragenic if they overlapped and were on the same strand as a protein-coding gene, long non-coding RNA, or short non-coding RNA other than the miRNA gene of interest, and as intergenic otherwise. Gene coordinates and strand information was obtained from Ensembl. MiRNA clusters (miRNA genes with inter-miRNA distance < 10 Kb) were copied from the miRBase website. Any precursor that was absent from the miRBase clusters was assigned to its own cluster.

#### 4.7.3 Colocalization with host gene eQTLs

eQTL summary statistics were generated using genotype data and post-mortem dlPFC transcriptomic profiles from 621 ROS/MAP participants as described in ref^21^. Correlation between miRNA and transcript abundance was estimated using Spearman correlation after regressing out batch, PMI, RIN, sex, age at death, study, clinical diagnosis of AD, and SVs from normalized miRNA data and regressing out batch, PMI, RIN, sex, age at death, and clinical diagnosis and SVs from normalized transcript data.

For intragenic miRNAs, we characterized colocalization between miR-QTLs and host gene eQTLs using R package ‘coloc’. We first applied the SuSiE wrapper to identify credible sets of causal variants for the miR-QTL and eQTL datasets separately. Then, following the recommendation in ref^77^, we tested for colocalization using a hybrid approach, allowing for multiple causal variants using function coloc.susie if credible sets were identified for both datasets, and assuming a single casual variant using function coloc.abf otherwise. Colocalization was defined as posterior probability of a shared causal variant (PP.H4) > 0.5.

#### 4.7.4 Brain pQTL and SMR analysis

To test whether the miR-QTLs were also pQTLs for predicted target proteins, we first gathered predicted targets for each eMiR using TargetScan 7.2 and RNA22 v2. For TargetScan, we downloaded context++ scores and selected targets with cumulative weighted context++ score < -0.2, as more negative scores indicate stronger targeting efficacy. For conserved miRNAs, we kept only targets with at least one conserved site. For RNA22 predictions, we used genes from our proteomics dataset and obtained the MANE transcript for each from Gencode. We then used the RNA22 batch submission program with the following parameters: sensitivity of 63%/specificity of 61%; seed size of 7 with 0 unpaired bases in seed; minimum 12 paired-up bases in heteroduplex; maximum folding energy of -12 Kcal/mol for heteroduplex; and no limit on number of G:U wobbles in seed region. We kept predictions with p-value < 0.05. We took the union of targets from the TargetScan and RNA22 predictions, with a median of 586 predicted target genes (interquartile range 291– 1053). We then used the --glm function from PLINK2 to test the effect of miR-QTL allele dosage on protein abundance of the predicted targets. The pQTL models included sex, protein SVs, and 10 genetic PCs as covariates. We performed multiple testing correction for the ∼17,400,000 SNP-protein tests and defined pQTLs at FDR < 5%.

For miR-QTLs that were also pQTLs for predicted target proteins, we used summary data-based Mendelian randomization (SMR^28^) to test whether the genetically regulated miRNA abundance mediates the effect of the miR-QTL on protein abundance. MiR-QTL and pQTL summary statistics were used as the exposure and outcome, respectively. SMR significance was defined as Bonferroni-corrected P-value < 0.05 using the number of SMR tests conducted. SMR association can arise from causality or pleiotropy of a single SNP that affects both the exposure and outcome, or from linkage of SNPs with distinct effects on the exposure and outcome. We used HEIDI^28^ to identify signals arising from causality or pleiotropy (HEIDI P-value ≥ 0.05) rather than linkage (HEIDI P-value < 0.05).

#### 4.7.5 Promoter and enhancer enrichment

To test for enrichment of miR-QTLs in brain promoters and enhancers, the Cochran-Mantel-Haenszel test (function mantelhaen.test from R base package ‘stats’) was applied using three MAF strata: (0-0.1), [0.1-0.2), [0.2-0.5]. A 2×2 table was constructed within each MAF strata (i.e. ‘Is the SNP a miR-QTL?’ versus ‘does the SNP overlap a promoter?’). For enhancers, SNPs in the full test window (500 Kb symmetric padding around miRNA gene) were considered. For promoters, SNPs within 50 Kb of miRNA precursors that were tested in our study were considered. This window was chosen based on the expected proximity of promoters to the gene and given that miRNAs may be found in clusters or within host genes^78^.

#### 4.7.6 Overlap with miR-QTLs from fetal neocortex and blood

The SNPs in the fetal neocortex miR-QTL study^34^ were reported by genomic position with respect to human reference GRCh38. To match them with our results, we applied LiftOver to convert SNP positions to GRCh37 coordinates. SNPs in the blood miR-QTL study were matched based on rsID.

For each miRNA with a miR-QTL in the discovery set (i.e. the fetal neocortex set or the blood set), we selected the miR-QTL with the lowest P-value and looked up its P-value, if available, in the replication set (our adult dlPFC set). We used these P-values to calculate the *π*_1_statistic (representing the fraction of true positives) as 1-*π*_0_, where *π*_0_ is the fraction of true negatives estimated using function ‘qvalue’ from the R package qvalue.

#### 4.7.7 Identification of brain miRNAs consistent with a causal role

We integrated human brain genome-wide miRNA profiles with results from GWAS of brain traits using multiple independent but complementary approaches. First, we performed a miRNA-wide association study (miRWAS) using FUSION^35^, which consisted of estimating miRNA weights from the reference ROS/MAP dataset including both genetic and miRNA expression data. Next, the miRNA weights were integrated with the GWAS summary statistics of neurodegenerative and psychiatric traits. For the weight calculation, we restricted ROS/MAP genome-wide genotyping to an LD reference panel of 1,190,321 SNPs provided by FUSION. Next, the SNP-based heritability for each miRNA was estimated using the miRNA and genetic data. For miRNAs with heritability p-value <0.01, FUSION computed the effect of SNPs on miRNA abundance using multiple predictive models (top1, blup, lasso, enet, bslmm), and the most predictive model was selected. For each miRNA, SNPs were selected within a 500Kb window around the miRNA. Finally, the genetic effects, or GWAS Z-scores, of neurodegenerative or psychiatric traits were combined with miRNA weights by calculating the linear sum of Z-score × weight to perform miRWAS.

Next, we determined whether the identified miRNAs and the brain trait share causal variants through colocalization analysis using coloc^20^. FUSION includes an interface to the coloc software to compute an approximate colocalization statistic based on the marginal FUSION weights^35^. We used coloc to estimate the posterior probability that the miRNA and traits share a causal variant (PP.H4).

Subsequently, we performed SMR^28^ to verify that the identified miRNAs mediated the association between the GWAS loci and the brain trait and to rule out association due to linkage disequilibrium with the HEIDI test^28^.

In summary, a miRNA was declared as being consistent with a causal role in a brain trait if it met all of the following four criteria: i) the miRNA was associated with the brain trait in the miRWAS at FDR P-value < 0.05; ii) the identified miRNA and brain trait share a causal variant as reflected by the posterior probability of a shared causal variant (PP.H4) >0.5 per coloc); iii) the miRNA showed evidence for mediating the association between the GWAS loci and the brain trait in the Mendelian randomization analysis at SMR P-value < 0.05, and iv) the mediation was not due to linkage disequilibrium based on HEIDI^28^ (HEIDI P-value ≥ 0.05).

#### 4.7.8 Mediation analysis of multi-omics signals in the same loci

Among the 4 identified causal miRNAs in brain illnesses, two were regulated by miR-QTLs in proximity (±500Kb window) with eQTLs of causal mRNAs for the same brain illness, and one was regulated by miR-QTLs in proximity with pQTLs for a causal protein for the same brain trait (Supplementary Table 17). These causal mRNAs and proteins were from a recent published work^26^. Because of their proximity, we sought to determine whether the causal miRNAs mediated the association between miR-QTLs and causal mRNAs or protein, respectively, for a particular brain trait using SMR for two molecular traits^79^. We used miR-QTL summary data as the exposure, and eQTL or pQTL summary data as the outcome. Mediation was defined as SMR P-value <0.05 and HEIDI P-value ≥ 0.05.

## 5 Data availability

miRNA data and miR-QTL summary statistics will be available upon publication on synapse.org at <u>syn51247298</u>. ROS/MAP resources are under controlled access to protect patient privacy and can be requested at https://www.radc.rush.edu.

## 6 Online Resources

miRBase miRNA clusters https://www.mirbase.org/cgi-bin/mirna_summary.pl?org=hsa&cluster=10000, accessed Aug 22 2022

TargetScan Release 7.2 http://www.targetscan.org/cgi-bin/targetscan/data_download.vert72.cgi PyschEncode reference enhancer list http://resource.psychencode.org

## 7 Code Availability

The analyses were conducted using publicly available software and R packages. Conversion between genome builds was done using LiftOver (https://genome-store.ucsc.edu/). Adapter trimming of miRNA reads was done using Trimmomatic v0.36 (http://www.usadellab.org/cms/?page=trimmomatic). miRNA quantification was done using mirDeep2 v0.1.2 (https://github.com/rajewsky-lab/mirdeep2/). Genotype filtering and VCF manipulation was done using vcftools v0.1.13 (https://vcftools.github.io), bcftools v1.9 (https://github.com/samtools/bcftools), and PLINK v2.0a (https://www.cog-genomics.org/plink/2.0/).

Identification of population outliers was done using EIGENSTRAT (https://github.com/DReichLab/EIG). Linear regression to identify miR-QTLs was done using PLINK v2.0a. Clumping of miR-QTL signals was done using PLINK v1.90b53 (https://www.cog-genomics.org/plink/1.9/). Genomic interval overlaps were characterized using bedtools v2.27.0 (https://github.com/arq5x/bedtools2). Summary data-based Mendelian randomization was done using SMR v1.03 for miR-QTLs with target protein expression and v1.02 for miR-QTLs with GWAS traits (https://yanglab.westlake.edu.cn/software/smr/). miRWAS and colocalization analysis of miR-QTLs and GWAS traits was done using FUSION (including the FUSION interface to pass marginal FUSION weights to coloc) (http://gusevlab.org/projects/fusion/). Other analyses were done using R (version 3.6.0 unless otherwise noted) using base functions or the following R packages: data.table v1.14.2 for data manipulation; edgeR v.3.28.1 to normalize miRNA data; variancePartition v1.16.0 to estimate percent variance explained; sva v.3.34.0 for surrogate variable analysis; coloc v5.2.1 (using R version 4.1.1) for colocalization analysis of miR-QTLs and host gene eQTLs; qvalue v2.15.0 to calculate qvalues and replication statistics.

## Acknowledgements

We gratefully acknowledge the research volunteers and staff of ROS/MAP for their participation and contributions. This work was supported by I01 BX003853 (APW); IK4 BX005219 (APW); I01 BX005686 (APW); R01 AG056533 (APW, TSW); R01 AG075827 (APW, TSW); R01 AG072120 (APW, TSW). Work in the E.C.L. lab was supported by NIH grant and the MSK Core Grant P30-CA008748. ROSMAP is supported by NIA grants P30AG10161, P30AG72975, R01AG15819, R01AG17917, U01AG46152, and U01AG61356. This work was supported by the Emory University Emory Integrated Computational Core Facility (RRID:SCR_023525).

## Notes

https://www.synapse.org/#!Synapse:syn51247298

https://www.radc.rush.edu

### Competing Interest Statement

The authors have declared no competing interest.

