## Supplementary figures for "Genetic regulation of microRNAs in the older adult brain and their contribution to neuropsychiatric conditions"

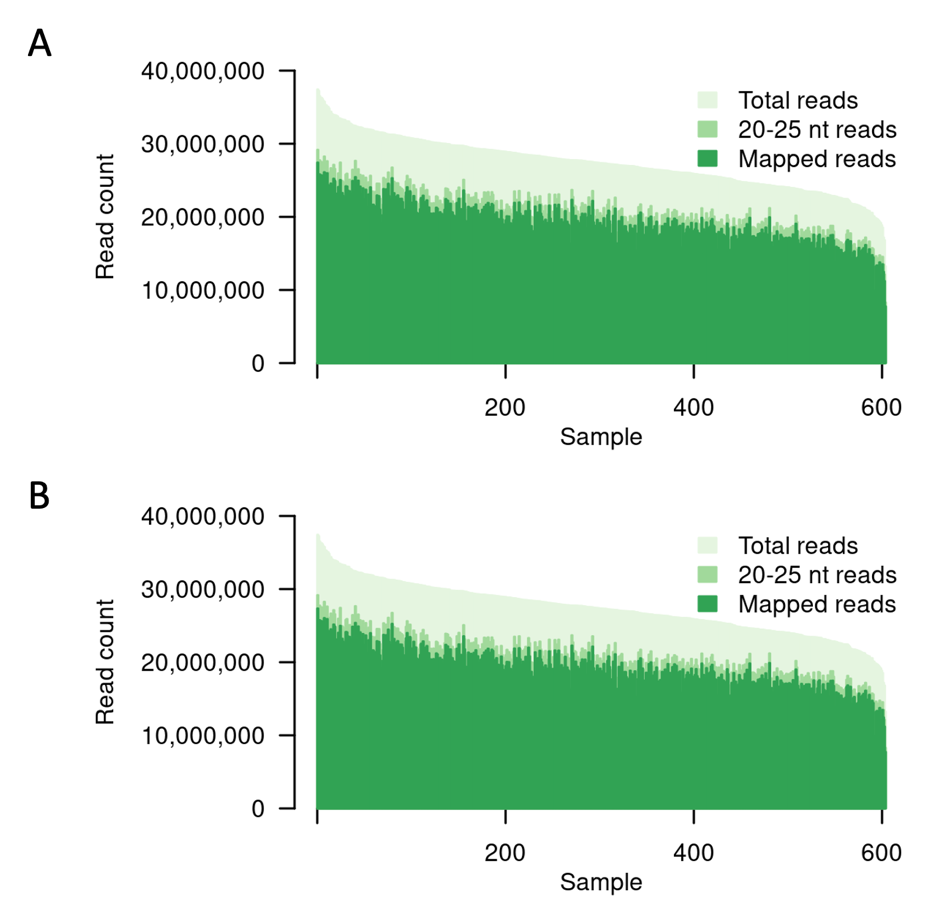

Supplementary Figure 1. Small RNA sequencing depth and mapped read counts. The count of total reads, reads with length 20-25 nt, and mapped reads are plotted in light green, medium green, and dark green, respectively. Samples are sorted by decreasing total read count. A) reads mapped to miRBase miRNAs; B) reads mapped to MirGeneDB miRNAs.

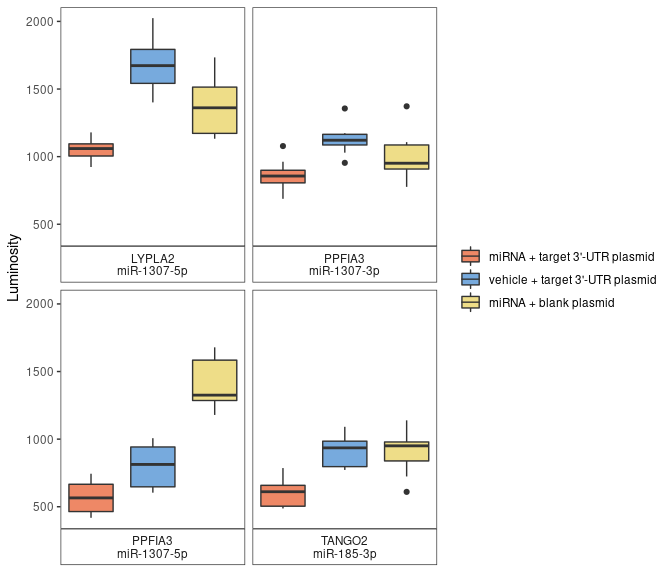

Supplementary Figure 2. Validation of miRNA-target interaction using luciferase assays. Each panel corresponds to one of the four miRNA-target pairs with significant SMR/HEIDI results. For all pairs, luciferase expression was significantly lower in assays transfected with the miRNA mimic than in assays transfected with vehicle only (red v. blue boxplots, p < 6×10^-4^ using linear regression with adjustment for plate effect), providing validation for miRNA-target interaction. The assays involving blank plasmid (yellow boxplots) were performed to verify that transfection with miRNA mimic did not interfere with luciferase readings. Each assay was performed in eight replicates. Center line indicates median, hinges indicate first and third quartiles, whiskers indicate 1.5 times interquartile range, and black points indicate observations outside that range.

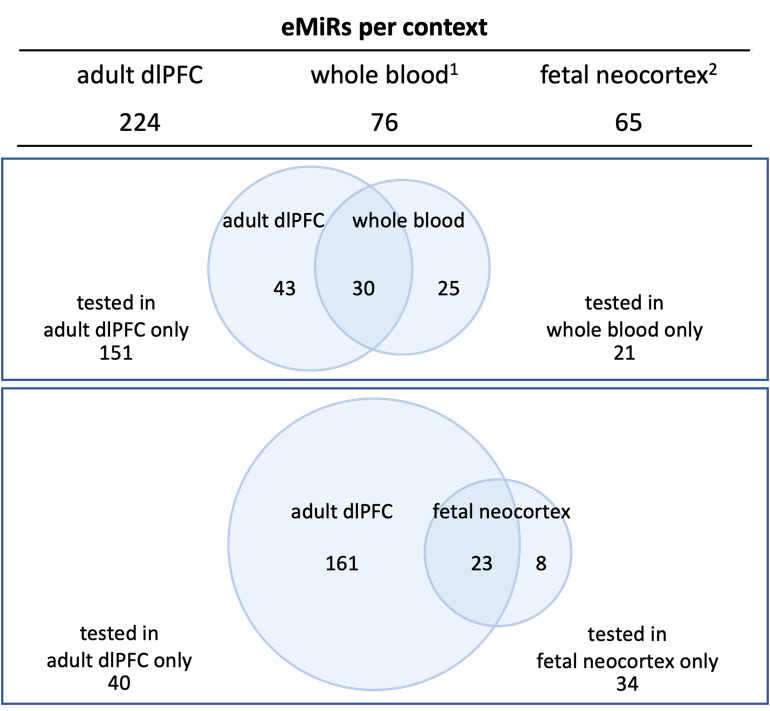

Supplementary Figure 3. Comparison of eMiRs (miRNAs with miR-QTLs) across contexts. The Venn diagrams summarize the number of shared and unique eMiRs between this adult dlPFC study and published eMiRs from whole blood and fetal (midgestation) neocortex. Seven miRNAs (miR-130b-3p/-5p, miR-204-5p, miR-31-5p, miR-323a-3p, miR-339-3p, miR-411-3p) were eMiRs in all three studies.

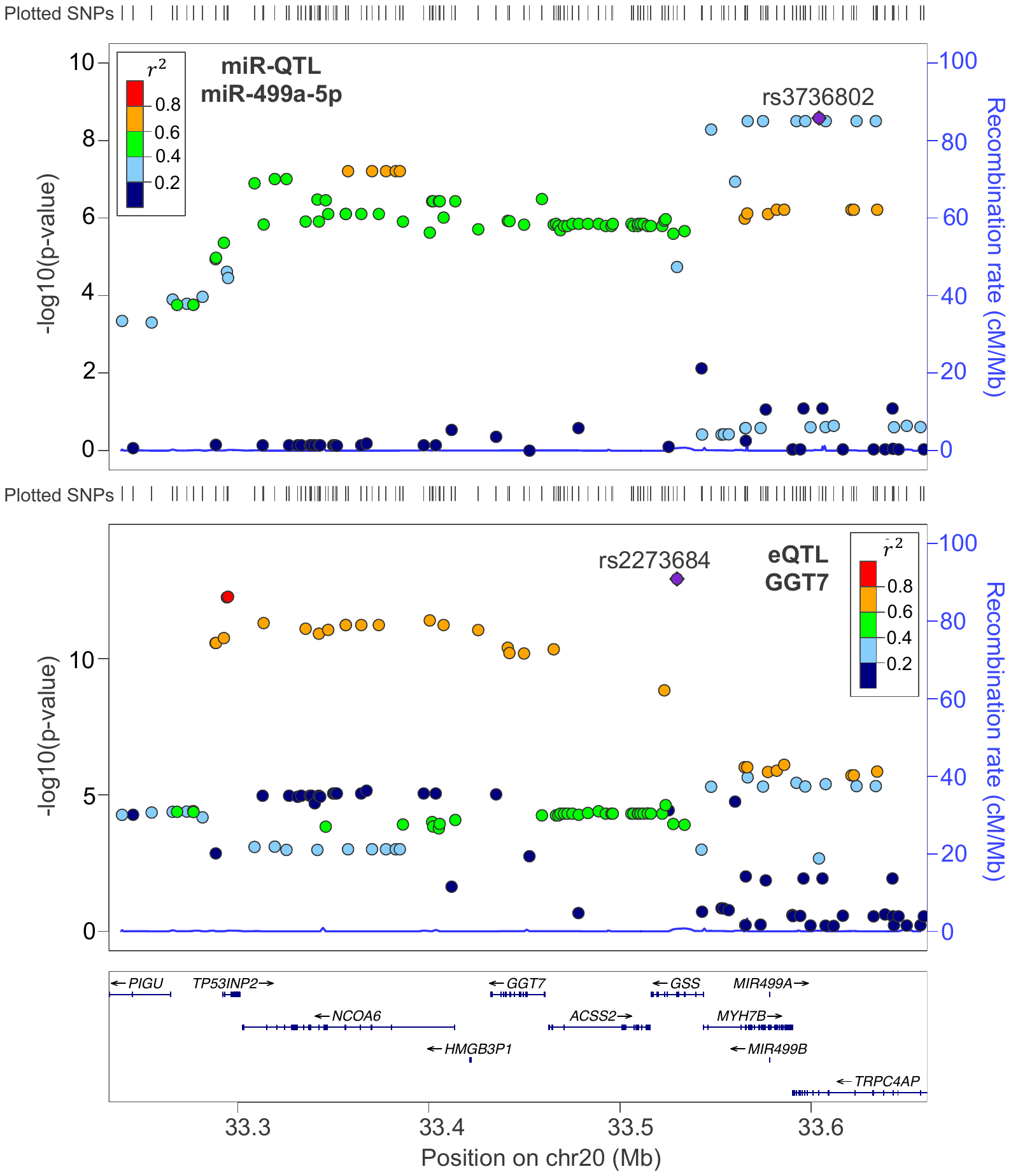

Supplementary Figure 4. LocusZoom plots for miR-499a-5p miR-QTL and GGT7 eQTL.

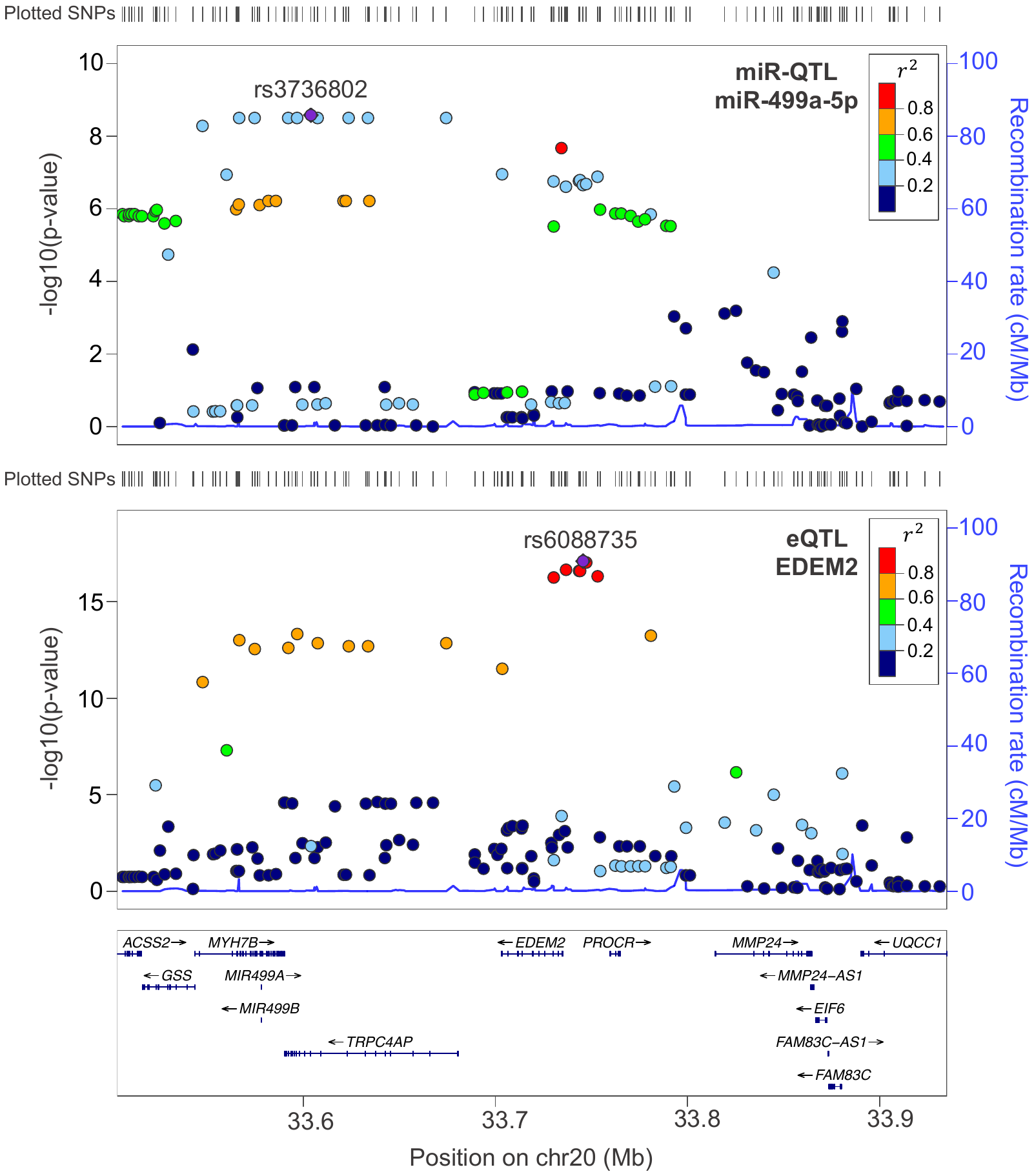

Supplementary Figure 5. LocusZoom plots for miR-499a-5p miR-QTL and EDEM2 eQTL.

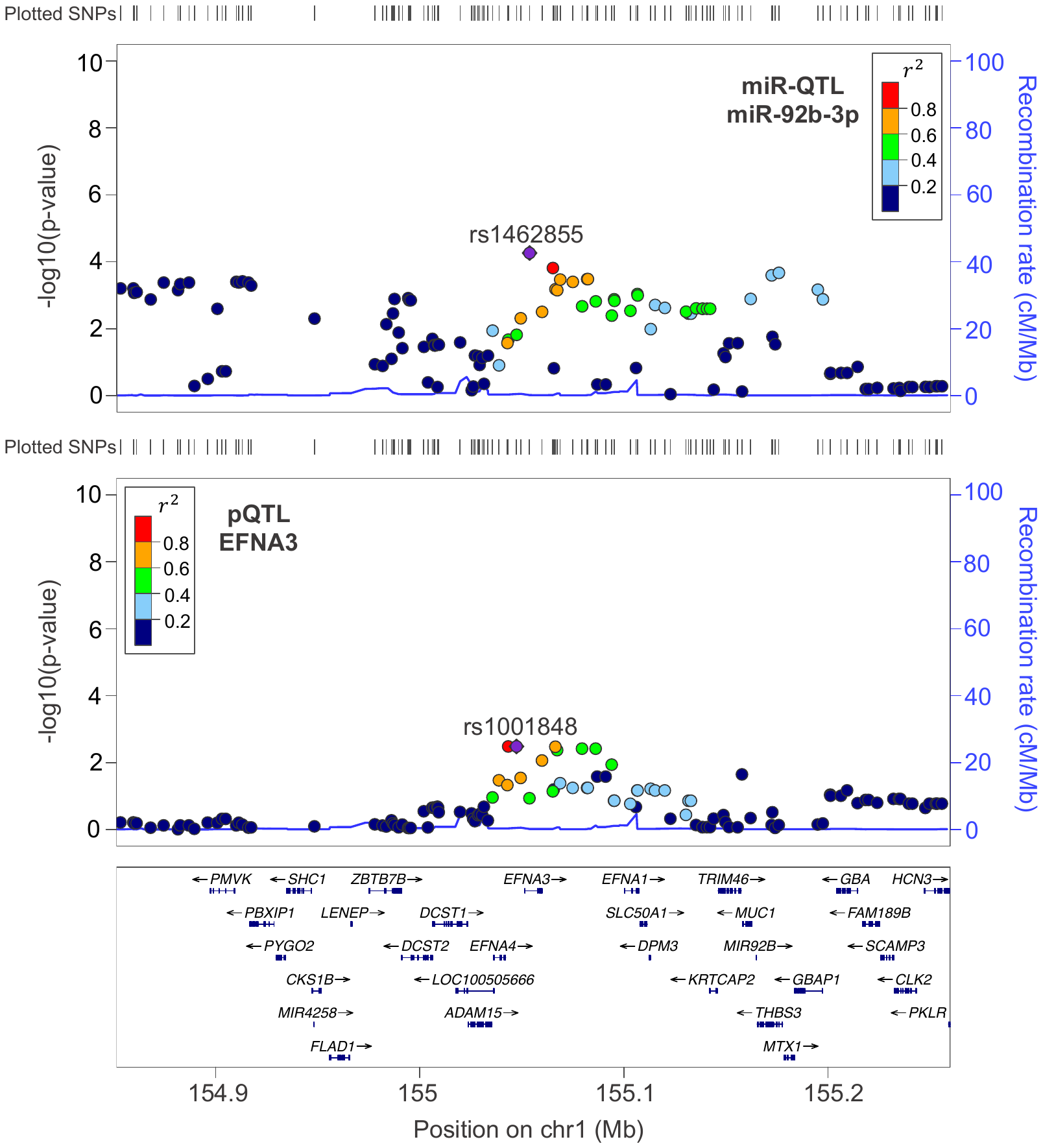

**Supplementary Figure 6.** LocusZoom plots for miR-92b-3p miR-QTL and EFNA3 pQTL.
